# Palmitoylation-dependent activation of NADK promotes NADP^+^ synthesis and tumorigenesis

**DOI:** 10.1101/2025.07.24.666230

**Authors:** Luojun Chen, Wanjun Zhang, Yue Zhu, Xiaoke Xing, Yi Yao, Huadong Pei, Yali Chen, Weijie Qin, Pingfeng Zhang

**Affiliations:** Renmin Hospital of Wuhan University, Wuhan, China; State Key Laboratory of Proteomics, National Center for Protein Sciences (Beijing), Beijing Institute of Lifeomics, Beijing, 100850, China; Department of Oncology, Georgetown Lombardi Comprehensive Cancer Center, Georgetown University Medical Center, Washington, DC, 20057, USA

## Abstract

NAD kinase (NADK) is the sole cytosolic enzyme that catalyzes the synthesis of nicotinamide adenine dinucleotide phosphate (NADP^+^) from NAD^+^. NADP^+^ is essential for anabolic reactions and redox balance. Here, we show that palmitate facilitates NADP^+^ synthesis by enhancing NADK palmitoylation and activity. NADK is post-translationally S-palmitoylated on three cysteine residues (Cys22, Cys23, and Cys26) within the amino-terminal domain by the protein-acyl transferase ZDHHC5, which stimulates NADK activity. Fatty acids activate NADK by enhancing its palmitoylation and relief of an autoinhibitory function inherent to its amino terminus. Furthermore, ZDHHC5^-/-^ mice showed defect in NADK palmitoylation and NADP^+^ production. Clinically, elevated expression of ZDHHC5 in pancreatic cancer patients was associated with increased NADK palmitoylation and correlated with poor prognoses for patients. These data reveal that fatty-acid and ZDHHC5-mediated palmitoylation has a critical role in NADP^+^ synthesis and cancers.

---

NADPH is a principal supplier of reducing power for anabolic reactions and redox balance, and its dysregulation leads to human diseases, such as aging^1^ and cancers^2,3^. Rapidly proliferating cancer cells require huge amount of NADPH as reducing agent for reductive synthesis of nucleic acids, proteins, and lipids^2^. NADK is the sole cytosolic enzyme that catalyzes the *de novo* formation of NADP^+^ by phosphorylation of NAD^+^^4,5^. NADK plays a crucial role in cell metabolism, survival, and oxidative defense in a variety of organisms^6,7^. Mammalian NADK has evolved an additional N-terminal domain compared to bacteria NADK, which helps mammalians to adapt to a variety of environments ^4,8^. Several regulatory mechanisms on this N-terminal domain have been discovered in recent years, including phosphorylation by PI3K/AKT ^9,10^ or PKC ^11,12^. Akt-mediated phosphorylation of NADK stimulates its activity to increase NADP^+^ production through relief of an autoinhibitory function inherent to its amino terminus.

Metabolic reprogramming is a hallmark of cancer cells to allow adaptation of cells to sustain survival signals^13,14^. Dysregulation in lipid metabolism is among the most prominent metabolic alterations in cancers^15,16^. 16-C saturated fatty acid palmitate could be attached to a cysteine residue via a thioester linkage^17^. Protein S-palmitoylation is a reversible, enzymatic posttranslational modification of proteins, which regulates protein functions by affecting their association with membranes, protein-protein interactions, trafficking, and stability^18–23^. Protein S-palmitoylation is catalyzed by 23 mammalian palmitoyl-transferases—known as DHHCs because of the conserved Asp-His-His-Cys sequence motif and is removed by acyl protein depalmitoylases^24^. Although more and more evidence supports the role of palmitoylation in cancers, their roles in NAD metabolism is still elusive.

NADK is the sole cytosolic enzyme that catalyzes the *de novo* synthesis of NADP^+^ ^6^. We first inquired whether palmitic acid (PA) induced S-palmitoylation would affect NADK activity. Using a click-chemistry-reaction (CCR)-based method and an acyl-biotin exchange (ABE) assay, we found that NADK was palmitoylated in HEK293T cells (Fig. 1a, b). We performed further studies using multiple pancreatic cancer cell lines and confirmed that ectopically expressed and endogenous NADK is S-palmitoylated through thioester bonds that are cleavable by treatment with hydroxylamine (NH2OH) (Extended Data Fig.1a, b). PA induced palmitoylation of NADK in a dose dependent manner (Fig. 1c). The general protein palmitoylation inhibitor 2-bromopalmitate (2-BP) treatment abrogated NADK palmitoylation, further confirming this modification (Fig. 1d and Extended Data Fig.1b).

**Figure 1.**
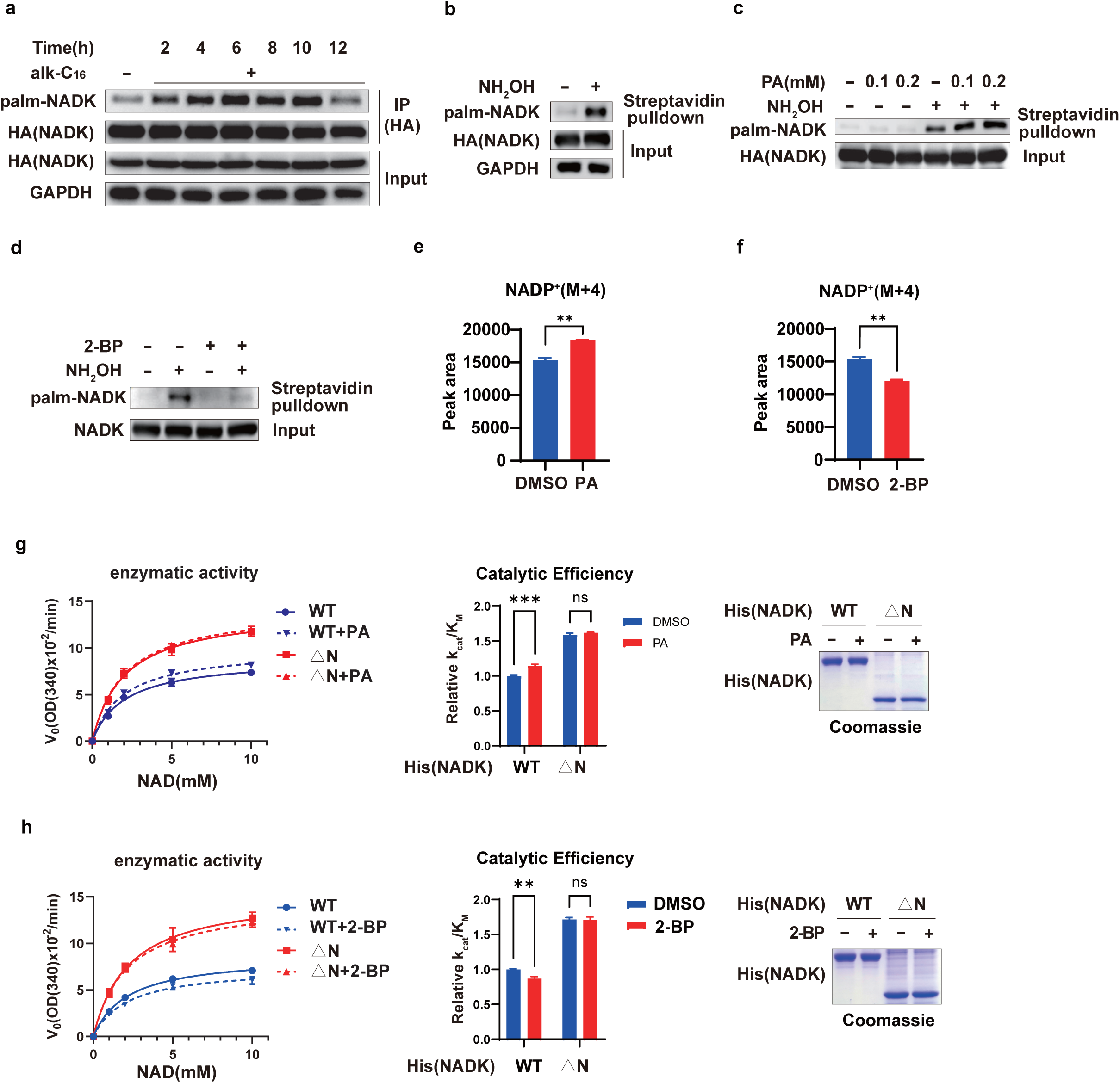
Palmitate induced palmitoylation promotes NADP^+^ synthesis. **a.** HEK293T cells were transfected with HA-NADK plasmid and labelled with 50 μM alk-C16 for the indicated time. Immunopurified NADK were subjected to CCR method and the NADK palmitoylation level was determined with the anti-biotin antibody. b. HEK293T cells were transfected with HA-NADK, and NADK palmitoylation was analyzed by ABE assay. c. HEK293T cells were treated with or without PA (0 ∼ 0.2 mM) for 12 h, and NADK palmitoylation was analyzed by ABE assay. d. HEK293T cells were treated with or without 0.1 mM 2-BP for 12 h, and NADK palmitoylation was analyzed by ABE assay. e, f. HEK293T cells were treated with or without 0.1 mM PA (e) or 0.1 mM 2-BP (f) for 12 h, *de novo* synthesized NADP^+^ were measured by LC-MS. g, h. HEK293T cells were transfected with His-NADK plasmids and treated with or without 0.1mM PA (g) or 2-BP (h) for 12h. His-NADK proteins were purified using Ni sepharose, and NADK protein level and enzymatic activity was examined as described in the method. △N, NADK lacking the N-terminal region of amino acid 1-95. [e-h] Data are presented as the mean ± SD of biological duplicates and are representative of three independent experiments. *P* values were determined by two-way ANOVA. V0, reaction velocity; OD, optical density. \**P*<0.05, \*\**P*<0.01, \*\*\**P*<0.001.

To examine whether PA induced palmitoylation affects NADP^+^ synthesis, stable isotope tracing with ^13^C3-^15^N-nicotinamide was employed with a 1-hour pulse label^9,25^ (Extended Data Fig.1c). We treated HEK293T cells with 0.1 mM PA, then measured M+4 derivatives of NADP^+^ production using liquid-chromatography mass spectrometry (LC-MS). PA promoted NADP^+^ production (Fig. 1e), and 2-BP treatment decreased NADP^+^ synthesis in cells (Fig. 1f). Consistently, purified NADK protein from 293T cells stimulated with PA exhibited much higher enzymatic activity and catalytic efficiency (*k*cat*/K*M, where *k*cat is the rate of catalysis), compared to NADK from cells without PA (Fig.1g). Consistently, NADK protein from 293T cells with 2-BP treatment exhibited lower enzymatic activity and catalytic efficiency (Fig. 1h). These results indicate that PA induced palmitoylation boosts NADK enzymatic activity.

Protein S-palmitoylation is catalyzed by 23 palmitoyl-transferases (ZDHHCs) in human cells^23^. Co-expression of HA-NADK and Flag-ZDHHCx in HEK293T cells showed that ZDHHC3, 5, 8, 17 promoted NADK palmitoylation (Extended Data Fig. 2a). Reciprocal co-IP assay between ZDHHC3, 5, 8, 17 and HA-NADK showed that ZDHHC5 had the strongest interaction with NADK (Extended Data Fig. 2b, c). Using purified His-NADK to pulldown HA-ZDHHC5 in HEK293T cells further confirmed ZDHHC5-NADK interaction (Extended Data Fig. 2d).

Reciprocal co-IP assay between ZDHHC5 and NADK also supported this conclusion (Extended Data Fig. 2e). Furthermore, overexpression of ZDHHC5 increased NADK palmitoylation in a dose dependent manner, and ZDHHC5 enzyme deficient mutant C134S did not (Fig. 2a). Knockout of ZDHHC5 in HEK293T cells abrogated NADK palmitoylation (Fig. 2b). These results validated ZDHHC5 is the major palmitoyl-transferase for NADK.

**Figure 2.**
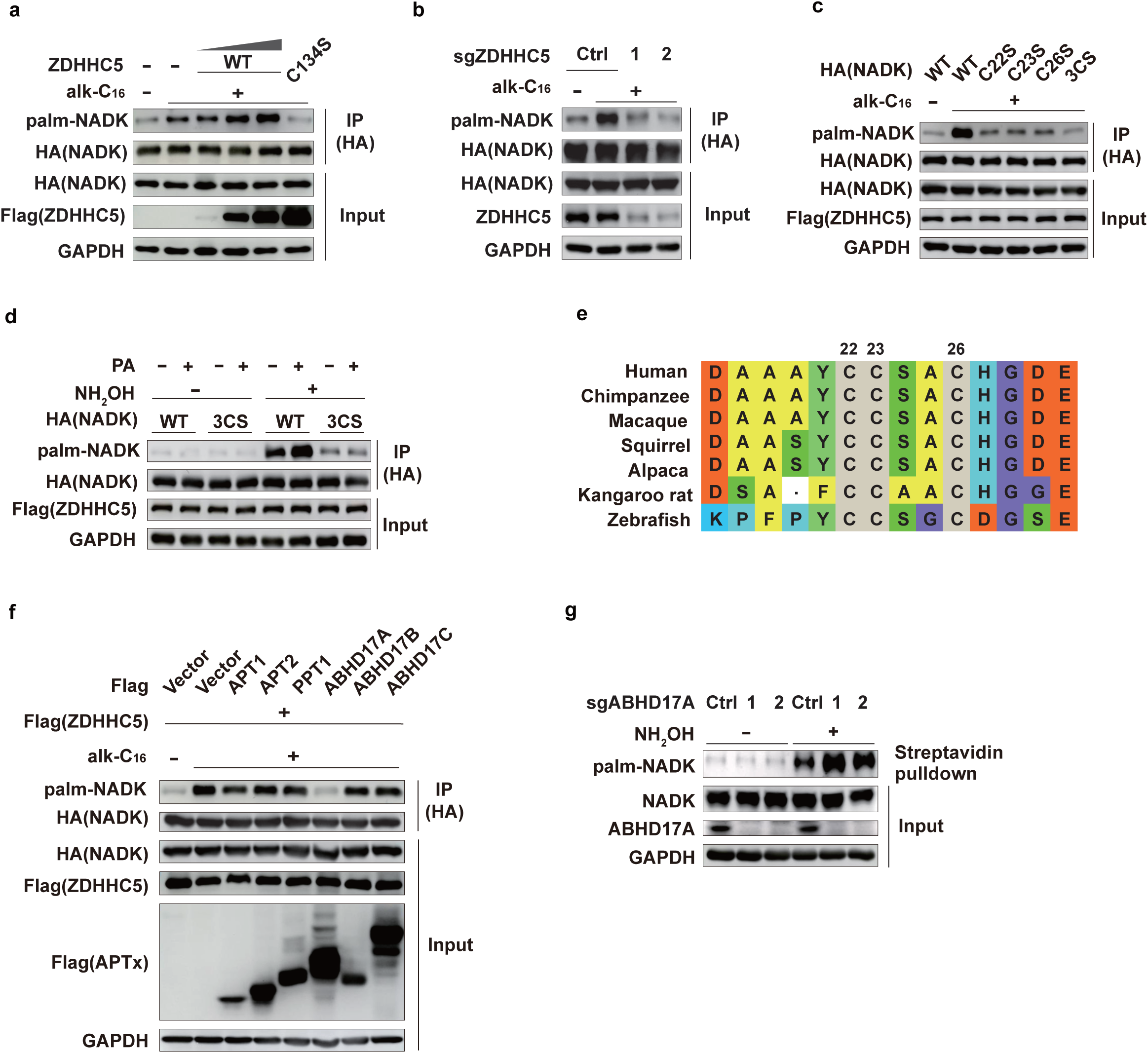
NADK is palmitoylated by ZDHHC5 at Cys22, 23 and 26. a. HEK293T cells were transfected with HA-NADK and gradient Flag-ZDHHC5 WT or C134S mutant and labelled with 50 μM alk-C16 for 8 h. NADK palmitoylation level was determined by CCR method. b. Endogenous ZDHHC5 was depleted in HEK293T cells by ZD5sgRNA1 and ZD5sgRNA2, followed by transfection with HA-NADK plasmid, and NADK palmitoylation level was determined by CCR method. c. As in b, various NADK cysteine mutants (C22S, C23S, C26S, or 3CS) were used for palmitoylation detection by CCR method. d. Endogenous NADK was depleted in HEK293T cells by sgRNA, followed by stable reconstitution with cDNAs encoding HA-NADK WT or 3CS mutant, and then cells were transfected with Flag-ZDHHC5 and treated with or without 0.1 mM PA for 12 h. NADK palmitoylation was analyzed by ABE assay. e. Sequence alignments of the consensus palmitoylation motif in NADK from human and other species. f. Endogenous NADK was depleted in HEK293T cells by sgRNA, followed by stable reconstitution with cDNA encoding HA-NADK WT, and then co-transfected with Flag-ZDHHC5 and Flag-depalmitoylases (APT1, APT2, PPT1, ABHD17A/B/C), these cells were metabolic labelled with 50 μM alk-C16 for 8 h, and NADK palmitoylation level was determined by CCR method. g. Endogenous ABHD17A was depleted in HEK293T cells by sgRNAs, then palmitoylation levels on endogenous NADK were determined by ABE method.

To identify which cysteine residue involves palmitoylation, we mutated each cysteine residue in NADK to serine and performed CCR assay. Mutations on cysteine 22, 23, 26 significantly decreased NADK palmitoylation (Extended Data Fig. 2f). We also generated a 3CS mutant with C22S/C23S/C26S, which almost abrogated NADK palmitoylation (Fig. 2c), indicating that these three cysteine residues are the major palmitoylation sites of NADK. Moreover, PA increased palmitoylation on WT NADK, but not 3CS mutant (Fig. 2d). 2-BP diminished palmitoylation on WT NADK, but not 3CS mutant (Extended Data Fig. 2g). NADK C22, C23, and C26 are conserved from fish to human (Fig. 2e), which suggests palmitoylation mediated regulation on NADK activity is evolutionarily conserved.

Protein palmitoylation is reversed by depalmitoylases. The palmitoylation-depalmitoylation cycle controls protein functions^26^. Overexpression of six protein depalmitoylases in HEK293T cells showed that ABHD17A significantly decreased NADK palmitoylation (Fig. 2f). Consistently, silencing ABHD17A increased NADK palmitoylation (Fig. 2g). Reciprocal co-IP assay and pulldown assay confirmed a specific interaction between ABHD17A and NADK (Extended Data Fig. 2h-i). Taken together, ABHD17A is the major depalmitoylase for NADK.

As the rate limiting enzyme to produce NADP^+^, knockout of NADK decreased NADP(H) level significantly (Extended Data Fig. 3a). Knockout of ZDHHC5 also decreased *de novo* synthesis of NADP(H) (Fig. 3a). To investigate the functional significance of NADK palmitoylation, we generated NADK knockout cells by Crispr/Cas9, and reconstituted these cells with gRNA-resistant WT NADK or the 3CS mutant. WT NADK cells rescued NADP(H) level, and 3CS mutant only partially rescued the NADP(H) level to about 70% to that of WT cells (Extended Data Fig. 3b). ^13^C3-^15^N-nicotinamide tracing also demonstrated that 3CS mutant had diminished ability to synthesize NADP(H) compared with WT NADK (Fig. 3b). Consistently, WT NADK purified from cells had higher enzymatic activity and catalytic efficiency than 3CS mutant (Extended Data Fig. 3c). But WT NADK purified from *E. coli* cells had similar enzymatic activity and catalytic efficiency to 3CS mutant (Extended Data Fig. 3d). In *E. coli* cells, NADK wasn’t palmitoylated, so 3CS mutant had no effect on the basal activity of NADK.

**Figure 3.**
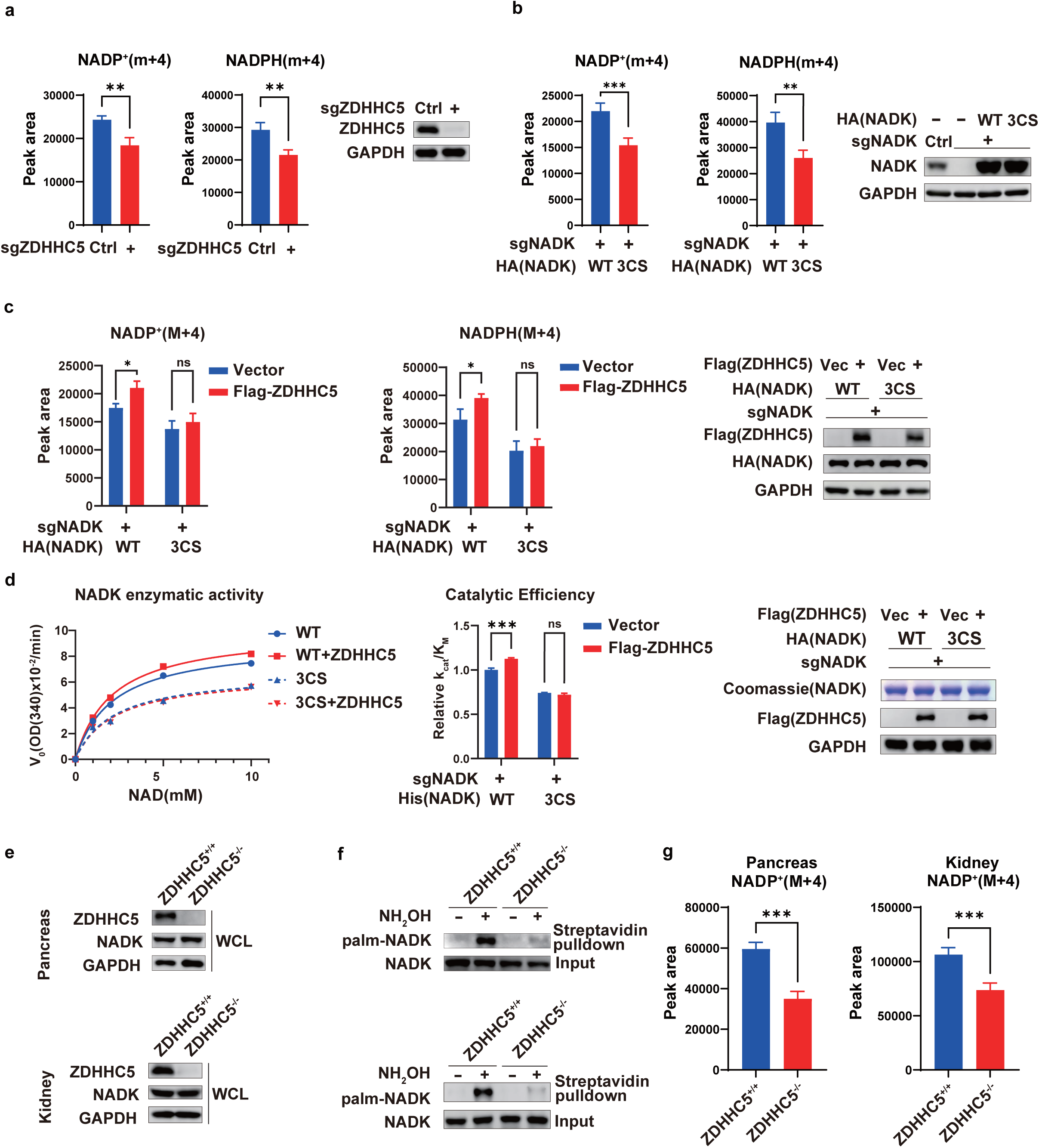
ZDHHC5 mediated NADK palmitoylation promoted NADK activity and NADP^+^ synthesis *in vitro* and in mice. a. Endogenous ZDHHC5 was depleted in HEK293T cells by sgRNA, and *de novo* synthesized NADP(H) measured by LC-MS (normalized peak areas) are presented. b. Endogenous NADK was depleted in HEK293T cells by sgRNA, followed by stable reconstitution with cDNAs encoding HA-NADK WT or 3CS mutant. NADK expression was immunoblotted and *de novo* synthesized NADP(H) was measured and presented as in a. c. As in b., but the cells were transfected with Flag-ZDHHC5. [a-c] Data are presented as the mean ± SD of biological duplicates and are representative of three independent experiments. [a-b] *P* values were determined by unpaired, two-tailed Student’s *t*-tests. [c] *P* values were determined by two-way ANOVA. \**P*<0.05, \*\**P*<0.01, \*\*\**P*<0.001. d. Similar as in c, the cells were stably reconstituted with cDNAs encoding His-NADK WT or 3CS mutant, and then were transfected with Flag-ZDHHC5. The NADK proteins were purified using Ni sepharose, and subjected to quantification and NADK activity assay. Data are presented as the mean ± SD of biological duplicates and are representative of three independent experiments. *P* values were determined by two-way ANOVA. V0, reaction velocity; OD, optical density. e, f. Tamoxifen induction applied to 6-8 weeks old ZDHHC5^flox/flox^ and ZDHHC5^flox/flox^/CreER^TM^ mice, ZDHHC5 expression was detected in the pancreas and kidney (e), subsequently, the NADK palmitoylation in pancreatic tissue or kidney were analyzed by ABE assay (f). g. NADP(H) level in pancreas or kidney from tamoxifen induced 6-8 weeks old ZDHHC5^flox/flox^/CreER^TM^ and ZDHHC5^flox/flox^ mice were analyzed by LC-MS (normalized peak areas). Data are presented as the mean ± SD of biological duplicates and are representative of four independent experiments. *P* values were determined by unpaired, two-tailed Student’s *t*-tests. \*\*\**P*<0.001.

ZDHHC5 overexpression stimulated NADP(H) synthesis in cells expressing wild-type NADK but failed to do so in cells with 3CS mutant (Fig. 3c). This result implies that ZDHHC5 regulates NADK activity via palmitoylation on these three cysteine residues. Supporting this conclusion, overexpression of ZDHHC5 boosted WT NADK activity but not 3CS mutant (Fig. 3d). Similarly, PA or 2-BP treatment increased or decreased WT NADK activity respectively but had no effect on 3CS mutant NADK activity (Extended Data Fig. 3e, f). In addition, overexpression of ABHD17A decreased WT NADK activity significantly but did not affect 3CS NADK mutant (Extended Data Fig. 3g). Taken together, ZDHHC5 boosts NADP^+^ level in cell through palmitoylating NADK N-terminal. Previous studies showed that Akt-mediated phosphorylation of NADK stimulated its activity to increase NADP^+^ production through relief of an autoinhibitory function inherent to its amino terminus. Perhaps ZDHHC5 mediated palmitoylation of NADK functions in a similar mechanism.

To distinguish the effect of ZDHHC5 on NAD metabolism *in vivo*, we generated ZDHHC5 conditional knockout mice. The western blot confirmed ZDHHC5 was knocked out in both pancreas and kidney tissues (Fig. 3e). The results from ABE assay showed that NADK palmitoylation level significantly decreased in both pancreas and kidney tissues of ZDHHC5 CKO mice (Fig. 3f). This result supported ZDHHC5 mediated NADK palmitoylation *in vivo*. Furthermore, we performed [U-^13^C]-glucose tracing experiment in mice (*in vivo*). ZDHHC5 CKO mice diminished the ability to *de novo* synthesize NADP^+^ from NAD^+^, supporting the roles of ZDHHC5-NADK axis in NADP^+^ *de novo* synthesis *in vivo* (Fig. 3g).

Previous studies showed that AKT boosts NADK activity through phosphorylating N-terminal region of NADK ^9^. We therefore investigated whether ZDHHC5-mediated NADK palmitoylation crosstalks with AKT-dependent phosphorylation. PA treatment didn’t affect NADK phosphorylation by AKT (Extended Data Fig. 4a). NADK palmitoylation deficient mutant also showed similar phosphorylation to wild type NADK (Extended Data Fig. 4b). These results indicate that palmitoylation does not interfere with AKT-mediated phosphorylation. On the other hand, insulin treatment or NADK phosphorylation mutant didn’t affect NADK palmitoylation (Extended Data Fig. 4c, d). NADK palmitoylation deficient mutant also didn’t affect G6PD-NADK interaction (Extended Data Fig. 4e), which is important for NADPH production ^27,28^. In conclusion, ZDHHC5-mediated NADK palmitoylation functions in parallel with the phosphoinositide 3-kinase (PI3K)-Akt pathway.

NADPH is a key co-factor in proliferating cells where it provides reducing power for biosynthesis, and its concentration in cell is controlled by NADK ^29,30^. NADK activity is upregulated in cancer cells, but the underlying mechanism is still elusive ^30–32^. We set out to explore the effect of palmitoylation on NADK in cancers. 2-BP treatment or ZDHHC5 knockout in pancreatic cancer cells MIA paca2 and PANC1 reduced cell proliferation and colony formation (Extended Data Fig. 5a-d). Endogenous NADK depletion in MIA paca2 and PANC1 cells leads to decreased colony formation (Extended Data Fig. 5e, f). Then cDNAs encoding WT and 3CS mutant NADK was stably reconstituted into NADK knock out cells for tumor growth studies. The exogenous proteins were expressed at similar levels but had elevated abundance compared with endogenous NADK (Extended Data Fig. 5e, f). The 3CS mutant had diminished colony formation compared with WT NADK (Extended Data Fig. 5e, f). To test the *in vivo* relevance of these findings in the context of cancer, we conducted xenograft studies with PANC1 cells implanted in nude mice. NADK knockdown significantly reduced tumor growth (Fig.4a and Extended Data Fig. 5g). The 3CS mutant had reduced tumor growth compared with WT NADK (Fig. 4a and Extended Data Fig. 5g), indicating that ZDHHC5 mediated NADK palmitoylation promotes tumor growth *in vivo*.

**Figure 4.**
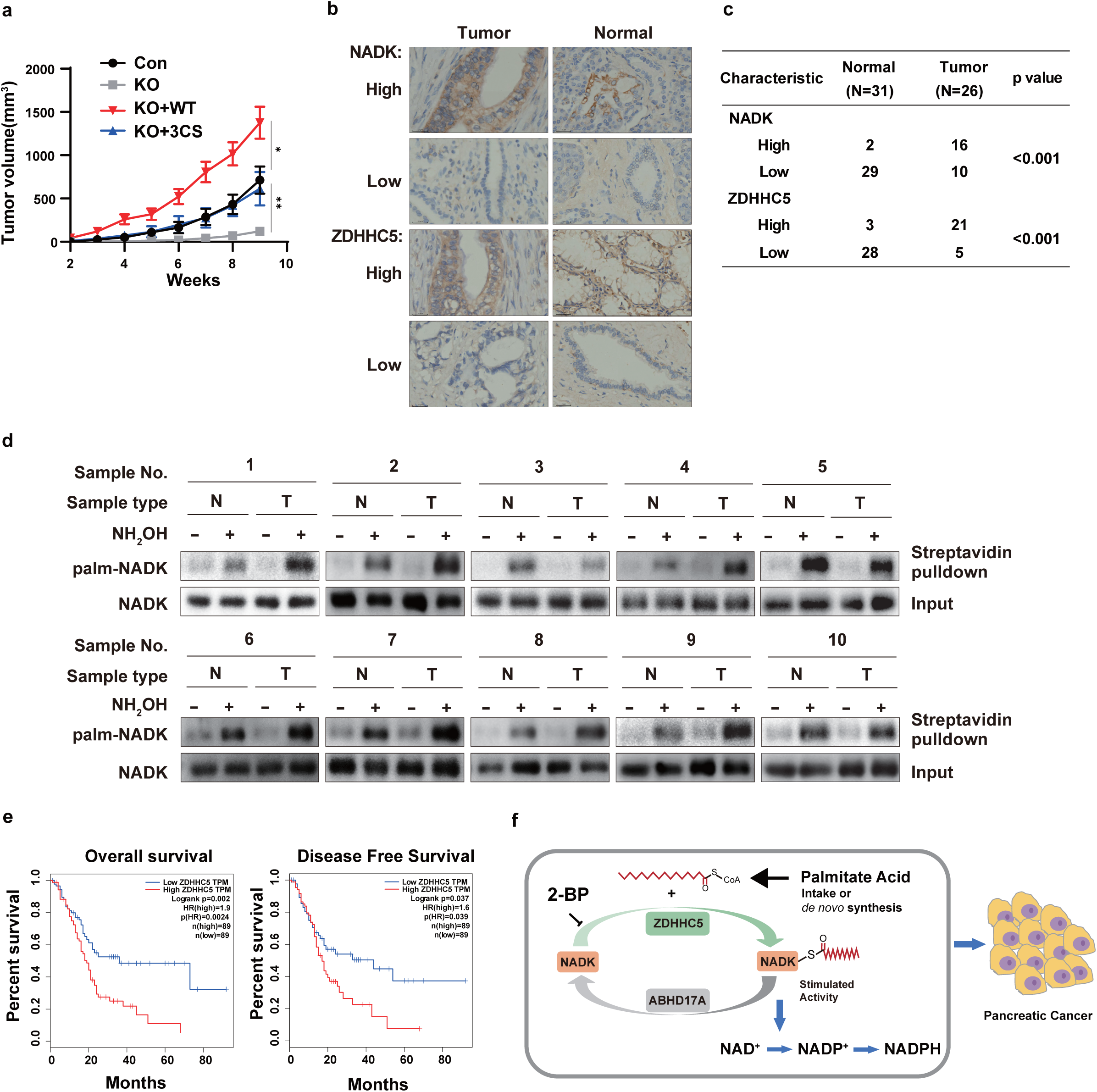
ZDHHC5-NADK axis is upregulated in PDAC. a. NADK knockout PANC-1 cells were stably reconstituted with NADK WT or 3CS mutant or control vector, then subcutaneously injected into athymic nude mice. Growth curve of the tumors over time are shown, error bars represent means ±SEM of 5 tumors. *P* values were determined by unpaired, two-tailed Student’s *t*-tests. b, c. Immunohistochemical staining for ZDHHC5 and NADK in PDAC and pancreatic normal tissues (b). Scale: 50μm. Chi-Squared Test for NADK and ZDHHC5 protein expression in PDAC tumor and normal pancreatic samples (c). d. The palmitoylation of NADK in 10 tissue samples of PDAC (T) and their paired pancreatic normal tissues (N) were analyzed by ABE assay. e. Kaplan-Meier curves for PDAC patients from TCGA databases, after stratification by the median level of ZDHHC5, were used for depicting overall survival and disease free survival. Patient data were analyzed using GEPIA (http://gepia.cancer-pku.cn). *P* values were determined by log-rank (Mantel-Cox) test. f. A working model for ZDHHC5 and ABHD17A mediated NADK palmitoylation stimulates NADP^+^ biosynthesis to promote pancreatic cancer.

Next, we examined the expression level of ZDHHC5 and NADK in cancers based on several public databases. Genotype-Tissue Expression (GTEx) database from GEPIA (http://gepia.cancer-pku.cn)^33^ showed that ZDHHC5 is elevated in several tumor types compared to normal tissue, including Lymphoid Neoplasm Diffuse Large B-cell Lymphoma (DLBC), Pancreatic adenocarcinoma (PAAD), Stomach adenocarcinoma (STAD), and Thymoma (THYM) ( Extended Data Fig.6a). ZDHHC5 mRNA level is significantly elevated in PDAC when comparing the PDAC tumor samples from The Cancer Genome Atlas (TCGA) database and pancreatic normal controls from the Genotype-Tissue Expression (GTEx) database (Extended Data Fig. 6b). Gene Expression Omnibus (GEO) database also showed ZDHHC5 mRNA level is significantly increased in PDAC compared to normal pancreatic tissue (Extended Data Fig. 6b, c). Moreover, mRNA levels in PDAC tumors and adjacent pancreatic normal tissues from the same patients (the GEO database) showed PDAC tumor has an elevated ZDHHC5 expression (Extended Data Fig. 6d, e). The Clinical Proteomic Tumor Analysis Consortium (CPTAC) database also showed that ZDHHC5 and NADK protein levels in PDAC tumor are much higher than that in the adjacent pancreatic normal tissues ^34^(Extended Data Fig. 6f, g). Together, ZDHHC5-NADK axis is upregulated in PDAC.

To further evaluate the roles of ZDHHC5-NADK axis in pancreatic cancers, we performed immunohistochemical staining of ZDHHC5 and NADK on the pancreatic cancer tissue samples, which were fresh-frozen samples obtained during surgical procedures. Upregulation of ZDHHC5 and NADK was detected in 81% (21 of 26) and 62% (16 of 26) of pancreatic cancer samples respectively, while 10% (3 of 31) and 6% (2 of 31) of adjusted normal pancreatic tissues showed positive staining for ZDHHC5 and NADK respectively (Fig. 4b, c, and Extended Data Fig. 7a-c). These results further support that both NADK and ZDHHC5 are upregulated in pancreatic cancers (Extended Data Fig. 7b, c). Indeed, ZDHHC5 is positively correlated with NADK expression in pancreatic cancer tissue, but not in pancreatic normal tissue (Extended Data Fig. 7d). Using ABE assay to examine NADK palmitoylation in pancreatic tumor tissues revealed that 70% tumor tissues had much higher NADK palmitoylation level (Fig. 4d). Furthermore, we examined the relationship between the ZDHHC5 expression and clinical outcome through GEPIA (http://gepia.cancer-pku.cn). The pancreatic cancer patients with low ZDHHC5 expression had significantly increased overall survival (OS) and progression-free survival (PFS) (Fig. 4e). We conclude that ZDHHC5 is upregulated in pancreatic cancer and ZDHHC5 mediated NADK palmitoylation may function as an independent prognostic predictor for pancreatic cancer patients.

Collectively, our results highlight a central role for ZDHHC5 mediated NADK palmitoylation in NADP^+^ *de novo* synthesis (Fig. 4f). This regulation links cellular nutrient, such as PA signaling to an acute increase in the production of cellular NADP^+^, which is the rate-limiting substrate for NADPH production. Our findings suggest that controlling NADK palmitoylation by down-regulation of ZDHHC5 or up-regulation of ABHD17A might be a potential clinical prevention strategy for NADPH associated diseases, such as pancreatic cancer.

## Materials and Methods

### Cell lines

HEK293T (ATCC, CRL11268), PANC-1, MIA PaCa-2 cells were cultured in Dulbecco’s Modified Eagle Medium (DMEM, Gibco) supplemented with 10% fetal bovine serum (FBS, Gibco), 100 U/ml penicillin and 100 μg/ml streptomycin and incubated at 37℃ with 5% CO2.

### Antibodies and other reagents

The antibodies that we used in this study are as follows: Rabbit monoclonal anti-Phospho-Akt (Ser473) (Cell Signaling Technology (CST), 4060), Rabbit monoclonal anti-Akt (CST, 4691), Rabbit polyclonal anti-NADK (CST, 55948), Rabbit monoclonal anti-p-RRXpS/pT (CST, 9624), Mouse monoclonal anti-Flag-M2 (Sigma, F1804), Mouse monoclonal anti-GAPDH (Proteintech, 60004-1-Ig), Mouse monoclonal anti-alpha Tubulin (Proteintech, 66031-1-Ig), Mouse monoclonal anti-HA (BioLegend, MMS-101P, Clone 16B12, RRID: AB_2314672), Mouse monoclonal anti-biotin (Abcam, ab201341), Rabbit polyclonal anti-ZDHHC5 (Sigma, HPA014670), Rabbit polyclonal antibody anti-ABHD17A (Thermo Scientific, PA5-46020). Rabbit polyclonal anti-ZDHHC5 (Proteintech, 21324-1-AP). Goat anti-mouse HRP (Jackson, 115035003 RRID: Ab_10015289), Goat anti-rabbit HRP (Jackson, 111035144 RRID: AB_2307391).

The critical reagents used in this study are in the following: Anti-Flag M2 Affinity Gel (Sigma, A2220), EZview Red Anti-HA Affinity Gel (Sigma, E6779), Streptavidin agarose beads (Thermo Fisher, SA10004), Ni sepharose 6Fast Flow (GE Healthcare, GE17-5318-01), Biotin picolyl azide (Click Chemistry Tools, 1167), Alkynyl palmitic acid (alk-C16) (Click Chemistry Tools, 1165), Tris(2-carboxyethyl) phosphine hydrochloride (TCEP) (Sigma, C4706), copper(II) sulfate (Sigma, 496130), Tris[(1-benzyl-1H-1,2,3-triazol-4-yl)methyl]amide (TBTA) (Sigma, 678937), palmitic acid (Sigma, P5585), N-ethylmaleimide (NEM) (Sigma, E3876), EZ-Link™ BMCC-Biotin (Thermo Scientific, 21900), 2-Bromohexadecanoic acid (Sigma, 21604), hydroxylamine (Sigma, 159417), EDTA-free protease inhibitors cocktail (Roche, 05892791001), Agarose (Biowest), Puromycin (Solarbo, P8230), Hygromycin B (Solarbo, IH0160), Imidazole (Solarbo, I8090), Thiazolyl Blue Tetrazolium Bromide solution (Beyotime, ST1537), β-nicotinamide adenine dinucleotide hydrate (NAD) (Sigma, N7004), D-glucose 6-phosphate disodium salt hydrate (G6P) (Sigma, G7250), glucose 6-phosphate dehydrogenase from baker’s yeast (G6PD) (Sigma, G6378), adenosine triphosphate salt (ATP) (Sigma, A2383), NADP/NADPH kit (Sigma, MAK038), IPTG (Sigma, I6758), ^13^C3-^15^N-nicotinamide (Cambridge Isotope Labs, CNLM-9757-0.001), [U-^13^C]-glucose (Cambridge Isotope Laboratories, CLM-1396), Tamoxifen (Sigma, T5648), Corn Oil (Solarbo, IC9000), BCA protein assay kit (Tiangen, PA115), Polyethylenimine (PEI) (Polysciences, 23966-2), Polybrene (Santa Cruz, sc-134220), Benzonase (Vazyme, DD4301), PMSF (Solarbio, P8340), Nicotinamide-free DMEM (US biologicals, D9800-17), Thiamine Hydrochloride (B1) (Sigma, 47858), Folic acid (Sigma, F7876), Pyridoxal hydrochloride (Sigma, P6155), Riboflavin (Sigma, R9504), D-Glucose (Sigma, G7528), Sodium pyruvate (Sigma, P2256), 10 kDa spin column (Millipore, UFC5100).

### Generation of stable cell lines

Lentiviruses were produced as previously described (http://www.addgene.org/protocols/plko/#E). Briefly, to generate stable NADK knockout, ZDHHC5 knockdown or knonckout, ABHD17A knockout, or NADK/ZDHHC5 double knockout cells, the following primers were used to target human NADK, ZDHHC5 or ABHD17A: sgNADK: 5′-TAGACTTCATCATCTGCCTG-3′; sgZDHHC5#1: 5′-CATCTGGCTTGCCAAAACGG-3′; sgZDHHC5#2: 5′-GGTGTGAACCCCTTCACCAA-3′; shZDHHC5#1: 5′-CGCACAACCAATGAACAGGTT-3′; shZDHHC5#2: 5′-CCAGTTACTAACTACGGAAAT-3′, sgABHD17A#1: 5′- ACAGCTGCTCACCTGGCACC-3′; sgABHD17A#2: 5′-GAGAAGAGGACCGTGTACCT-3′.

sgRNA were cloned into a lentiCRISPRv2 vector, and shRNA were cloned into a PLKO.1 vector. The retrovirus was produced by co-transfection of lentiCRISPRv2 or PLKO.1 with psPAX2 and pMD2.G plasmids. HEK293T, PANC-1, MIA PaCa-2 were infected with the retrovirus and selected with 4 ug/ml puromycin for 1 week. To obtain single cell clones, infected cells were diluted and seeded into 96-well plates. The knockout efficiency of NADK, ZDHHC5 or ABHD17A was confirmed by immunoblotting. To generate stable NADK rescue cells, HA-NADK, His-NADK or Flag-NADK (synonymous mutations were included in the region where the guide RNA sequence is targeting) were cloned into the retroviral pHAGE vector, then were co-transfected with pMD2.G and psPAX2 into NADK depleted HEK293T, PANC-1, or MIA PaCa-2 cells, following a selection with 300 μg/ml hygromycin B for 1 week.

### ABE assay

Briefly, cells or pancreatic cancer samples were lysed in lysis buffer (50 mM Tris-HCl, pH 7.4, 150 mM NaCl, 2% SDS) containing 1× EDTA-free protease inhibitors cocktail, 1 mM PMSF, 500 units/mL benzonase. The cell lysate was treated with 10 mM TCEP for 30 min with nutation at room temperature (RT) in the dark. NEM (1 M stock in ethanol) was added to a final concentration of 25 mM and incubated for 2 h at RT to block the free cysteine residues. Then terminated by methanol-chloroform-H2O (4:1.5:3) precipitation. The samples were re-suspended with 300 μL lysis buffer containing 5 mM EDTA, then divided into two groups, one group was treated with neutralized NH2OH at a final concentration of 0.75 M for 1 h at RT, the other group was added with equal volume of lysis buffer as a control. The reaction was terminated by methanol-chloroform-H2O (4:1.5:3) precipitation and resuspended in 150 μL lysis buffer. Next treated with 2 μM BMCC-biotin for 2 h at RT before a final methanol-chloroform-H2O precipitation. Then samples were re-suspended and immobilized on streptavidin agarose beads overnight. After washing three times with wash buffer (50 mM Tris-HCl, pH 7.4, 150 mM NaCl, 1% SDS), samples were boiled in 60 μL 2 × Laemmli sample buffer and subjected to immunoblotting with the indicated antibodies.

### Click chemistry reaction

Cells were metabolic labelled with 50 μM chemical reporter of alkynyl palmitic acid (alk-C16) for 8 h, washed twice with cold phosphate-buffered saline (PBS) and lysed in lysis buffer (50 mM triethanolamine (TEA), pH 7.4, 150 mM NaCl, 2% SDS, containing 1× EDTA-free protease inhibitors cocktail, 1 mM PMSF, and 500 units/mL benzonase). The supernatant was collected after centrifugation at 12,000 rpm for 10 min, then diluted with TEA buffer to 0.2% SDS. The target proteins were purified with anti-HA affinity gel, then eluted with 50 μL elution buffer (50 mM TEA, pH 7.4, 150 mM NaCl, 2% SDS), Click chemistry reagents were added to the eluted samples: 0.5 μL of 10 mM biotin azide, 0.5 μL of 10 mM Tris[(1-benzyl-1H-1,2,3-triazol-4-yl)methyl]amine (TBTA), 1 μL of 50 mM copper(II) sulfate, 1 μL of 50 mM Tris(2-carboxyethyl)phosphine hydrochloride (TCEP hydrochloride). The reaction mixtures were mixed thoroughly and incubated for 1 h in the dark at RT. Then, 15 μL of 5×Laemmli sample buffer was added and boiled at 95℃ for 10 min, finally analyzed by western blotting with anti-biotin antibody.

### Immunoblot analysis

After washing twice with cold PBS, cells were collected and lysed in NETN buffer (50 mM Tris-HCl, pH 8.0, 150 mM NaCl, 1% NP-40, 1 mM EDTA, containing 1 mM PMSF, 1 × EDTA-free protease inhibitors cocktail) on ice. After sonication, cell lysates were centrifuged at 12,000 rpm for 10 min at 4℃. The total protein was quantified using a BCA assay kit. About 10∼30 μg protein were analyzed by immunoblotting with indicated antibodies. The immunoblotting images were captured with GE Healthcare ImageQuant™ LAS 500.

### Immunoprecipitation and His-Tag Pull-Down assay

HEK293T cells were lysed in NETN buffer (50 mM Tris-HCl, pH 8.0, 150 mM NaCl, 1% NP-40, 1 mM EDTA, containing 1 mM PMSF, 1 × EDTA-free protease inhibitors cocktail), after gentle sonication, cell lysates were immunoprecipitated with anti-HA affinity gel or Anti-Flag M2 affinity gel at 4℃ overnight. After washing three times with lysis buffer, samples were subjected to immunoblotting with indicated antibodies.

pET-28a-NADK was transformed into *E. coli* BL21(DE3) and induced with IPTG to produce His-NADK recombinant protein. His-NADK protein was immobilized on Ni sepharose for 2 h at 4℃. HEK293T cells transfected with indicated plasmids were lysed in lysis buffer (50mM Tris-HCl, pH 7.4, 150mM NaCl, 1% NP-40, containing 1 mM PMSF, 1 × EDTA-free protease inhibitors cocktail), then incubated with Ni sepharose combined with his-NADK proteins at 4℃ overnight. After washing three times with lysis buffer, beads were boiled in 60 μL 2 × Laemmli sample buffer and subjected to immunoblotting with indicated antibodies.

### Protein purification

To purify His-NADK from HEK293T cells, endogenous NADK was knocked-out from HEK293T cells, then cDNAs encoding His-NADK WT or variants were stably reconstituted. Cells were lysed in lysis buffer (50 mM Tris-HCl, pH 7.4, 150mM NaCl, 1% NP-40, containing 1 mM PMSF, 1 × EDTA-free protease inhibitors cocktail) on ice, then centrifuged at 12,000 rpm for 10 min at 4℃, and supernatant were incubated with Ni sepharose at 4℃ overnight. After washing with wash buffer A (50 mM Tris-HCl, pH 7.4, 150 mM NaCl, 1% NP-40, containing 1 mM PMSF, 1 × EDTA-free protease inhibitors cocktail, 20 mM imidazole), His-NADK proteins were eluted using elution buffer (50 mM Tris-HCl, pH 7.4, 150 mM NaCl, 300 mM imidazole). Imidazole was removed using ultrafiltration centrifugal tube (10kD-cutoff) via buffer replacement with wash buffer B (50 mM Tris-HCl, pH 7.4, 150 mM NaCl).

For His-NADK purification from *E. coli* BL21(DE3), the bacterial culture was induced with 0.2 mM IPTG when OD600 reached 0.6, cells were harvested after growing at 18℃ overnight. The cell pellet was resuspended in ice-cold lysis buffer (50 mM Tris-HCl, pH 7.4, 150 mM NaCl, 0.1% Triton X-100, containing 1 mM PMSF, 1 × EDTA-free protease inhibitors cocktail). Cell lysates were centrifuged at 20000g for 20 min at 4℃, and the supernatants were incubated with Ni sepharose at 4℃ overnight. The following purification and imidazole removal was the same as the above description.

### NADK enzymatic assay

NADK activity was assayed as described (*Hoxhaj et al.*, 2019) with modifications. Approximately 0.5 μg of purified His-NADK WT or variants from HEK293T cells or *E. coli* were subjected to an NADK enzymatic assay that couples the generation of NADP^+^ to G6PD-mediated production of NADPH, which was measured as a change in the absorption at 340nm (A340) every 2 minutes for 20 min at 37℃ using Thermo Scientific™ Multiskan Sky Microplate Spectrophotometer. The assay was performed in a 100 μL reaction in 96-well plate containing 10 mM ATP, 10 mM glucose-6-phosphate, 0.5 U G6PD, 10 mM MgCl2, 100 mM Tris-HCl (pH 8.0), varying concentrations of the substrate NAD^+^ (0 mM, 1 mM, 2 mM, 5 mM, 10 mM), and His-NADK WT or variant proteins.

### Measurements of NADP^+^ and NADPH using a colorimetric enzyme-based assay

NADP(H) were measured using a modified NADP/NADPH colorimetric quantification assay (Sigma, MAK038) as previously described (*Hoxhaj G,* 2019). Briefly, metabolite extraction from a 10 cm culture dish per sample was performed on dry ice with 500 μL 80%(*v/v*) HPLC-grade methanol (pre-chilled at −80℃), incubated at −80℃ for 2h, then centrifuged at 14,000 g for 20 min at 4℃, and the supernatants were dried down under Speedvac/lyophilize. Dried metabolites were then resuspended in 200 μL of the manufacturer′s extraction buffer and centrifuged at 3000 g for 2 min. Supernatants from each sample were split in two halves. One half (A) was subjected to incubation at 60℃ for 30 min to decompose NADP^+^, leaving only NADPH. The other half (B), containing NADP^+^ and NADPH, was left on ice for 30 min. Split samples A and B were transferred to clear-bottom 96-well plates and the NADP^+^-cycling enzyme was added to each well for 5 min, resulting in conversion of NADP^+^ to NADPH in sample B, followed by addition of the manufacture′s NADPH developing solution. NADPH was continuously monitored at 450 nm every 15 min for 2 h. For normalization of each sample, the remaining pellet from the metabolite extractions were solubilized in 8 M urea in 10 mM Tris-HCl buffer (pH 8) and the total protein was quantified using a BCA assay. NADP^+^ concentrations are equal to the difference between sample B and A using the total protein.

### [^13^C3-^15^N]-nicotinamide tracing of *de novo* NADP^+^ generation in cells

[^13^C3-^15^N]-nicotinamide was used to trace the NADP(H) as described previously (*Hoxhaj et al.*, 2019). Briefly, cells were washed twice with PBS buffer and incubated in nicotinamide-free DMEM containing 4 mg/L of [^13^C3-^15^N]-nicotinamide and 10% FBS for 1 h. The nicotinamide-free DMEM containing [^13^C3-^15^N]-nicotinamide was compounded using nicotinamide-free DMEM supplemented with standard DMEM concentrations of Folic Acid, Pyridoxal, Riboflavin, Thiamine, Glucose, Sodium Pyruvate, and [^13^C3-^15^N]-nicotinamide, finally adjusted pH to 7.2. Then, cells were washed three with cold PBS buffer and metabolites were extracted on dry ice with 800 μL of 80%(*v/v*) HPLC-grade methanol (pre-chilled at −80℃), and incubated at −80℃ for 2h or overnight. The extraction was centrifuged at 20,000 g for 10 min at 4℃ and the supernatant was transferred to new centrifuge tube and dried down under Speedvac/lyophilize. The dried pellets were resuspended in HPLC-grade water and the samples were injected in ACQUITY UPLC H-Class system coupled with a 6500 plus QTrap mass spectrometer (AB SCIEX, USA) equipped with a heated electrospray ionization (HESI) probe. Extracts were separated by a synergi Hydro-RP column (2.0×100 mm, 2.5μm, phenomenex) with a flow rate of 0.5 mL/min. Eluent A was 2 mM triisobutylamine adjusted with 5 mM acetic acid in water, and eluent B was methanol. Separation was operated under the following linear or gradient conditions: 0∼1 min at 5% B; 1∼6 min, 5∼50% B; 6∼7 min, 98% B; 7∼8 min, 5% B. Column chamber and sample tray were held at 35℃ and 10℃. Multiple reaction monitor (MRM) mode were used to detect the metabolites. MS conditions were: nebulizer gas (Gas1), heater gas (Gas2), and curtain gas were set at 55, 55, and 35 psi, respectively. The ion spray voltage was −4500 V in negative ion mode. The optimal probe temperature was determined to be 500℃. Metabolite identification and peak integration were carried out by SCIEX OS 1.6 software.

### Mouse experiment

Procedures for mice experiments were performed in accordance with the guidelines of the Institutional Animal Care and Use Committee (IACUC) of National Center for Protein Sciences (Beijing). Mice were housed in a temperature-controlled environment with a 12-hour light/dark cycle, and fed ad libitum.

ZDHHC5^flox/flox^ mice was a gift from Dr. Dante Neculai (Zhejiang University). ZDHHC5^flox/flox^ mice were crossed with CAGGCre-ER^TM^ mice to obtain ZDHHC5^flox/flox^/CreER mice. All genetic models were on the C57BL/6 background. Both male and female mice (6-8 weeks old) were used for all described experiments. Mice genotypes were determined by PCR analysis on genomic DNA purified from mice tail tissue, and the genotyping primers are as follows: CreER forward: 5′-GCTAACCATGTTCATGCCTTC-3′, reverse: 5′-AGGCAAATTTTGGTGTACGG-3′; ZDHHC5 forward: 5′-TTAAGTTGCTTCAGAGATAGGAGTGTAAC-3′, reverse: 5′-TTGACACCAGCACAAATCTAAAGAG-3′.

### Tamoxifen-induced deletion of ZDHHC5 in mice

To achieve conditional knockout of ZDHHC5, 6-8 weeks old ZDHHC5^flox/flox^/CreER and ZDHHC5^flox/flox^ mice were intraperitoneal injected with tamoxifen (80 mg/kg body weight, dissolved in corn oil) for 7 successive days. After 7 days′ rest, mice were genotyped from tail to test the knockout efficiency.

### [U-^13^C]-glucose tracing isotope tracing of *de novo* NADP^+^ generation in mice

Mice were fasted 8 h before these experiments. Catheters (28-gauge) were placed in the lateral tail vein. [U-^13^C]-glucose infusions were started immediately after implantation of the catheter. The [U-^13^C]-glucose solution dissolved in 3 ml of saline was infused at a rate of 250 μL/h for 12 h, with a total dose of 14.4 g/kg. Animals were euthanized at the end of the infusion; then, pancreas and kidneys were dissected, rinsed briefly in cold saline, and flash frozen in liquid nitrogen.

For the extraction of NADP(H), 500 μL of 80% (*v/v*) HPLC-grade methanol (pre-chilled at −80℃) were added to 50 mg tissue. The tissue was smashed/ground for 1-2 min with tissue grinder at −30℃, and then incubated at −80℃ for 2 h or overnight. After centrifugation at 14,000g for 20 min at 4℃, the supernatant was transferred into a new tube to freeze dry. The metabolites were analyzed using LC-MS as describe in cells as above.

### Xenograft assays

Endogenous NADK-knockout PANC-1 cells stably reconstituted with cDNAs encoding HA-NADK WT or variants were suspended in PBS. Then 2×10^6^ cells (n=5 per group) were injected subcutaneously into BALB/c-nude mice (nu/nu, female, 5 weeks old). Tumors were measured (length and width) every two weeks by calipers to determine the tumor volume using the formula (0.5×length×width^2^).

### Soft agar colony formation assay

Cells (3000 cells/well for MIA PaCa-2 cells, 5000 cells/well for PANC-1 cells) were seeded in the top layer of 0.35% agarose premixed with DMEM supplemented with 10% FBS, 100 U/ml penicillin, and 100 μg/ml streptomycin in 6-well plates with a bottom layer of 0.7% agarose, and incubated at 37℃ with 5% CO2. Colonies were detected in 3 weeks for MIA PaCa-2 cells, 5 weeks for PANC-1 cells by staining with 0.5 mg/ml Thiazolyl Blue Tetrazolium Bromide solution for 3 h. Colony numbers were determined by OpenCFU software (http://opencfu.sourceforge.net).

### Tissue processing, immunohistochemistry and analysis

The paraffin sections of pancreatic cancer samples were obtained from the department of Pathology, Renmin Hospital of Wuhan University. All studies were approved by the Ethics Committee of the Renmin Hospital of Wuhan University, and informed consent was obtained from all patients. Formalin-fixed specimens were sliced continuously with a thickness of 4 μm, patched, and baked in an oven for 1 h. After dewaxing and hydration, paraffin sections were repaired in EDTA buffer in a microwave oven for 8 min, incubated in 3% H2O2 for 10 min, blocked with 5% BSA for 30 min at 37℃, and then incubated with anti-NADK (CST, 55948, 1:100), or anti ZDHHC5 (Proteintech, 21324-1-AP, 1:200) antibodies at 4℃ overnight in a humidified chamber, and were subsequently incubated with secondary antibodies, subjected to DAB coloration, hematoxylin staining, gradient ethanol dehydration, xylene transparent, and neutral gum sealing piece.

Images were acquired with a Digital pathology scanner (Leica^®^ Aperio Versa 8). Three areas were randomly selected for each image. The positively stained cells of each area were analyzed, and the mean staining intensity was calculated using Image-Pro Plus software. Paired two-tailed *t* test were used for difference analysis between the cancer and the adjacent tissue. The Pearson test was used for the analysis of the correlation between NADK and ZDHHC5.

### Sequence Alignments

Protein sequences of NADK from different species were obtained from Uniprot database (https://www.uniprot.org/), these NADK protein accession numbers were: O95544_HUMAN, K7CII1_PANTR, F7FZM8_MACMU, I3MJS9_ICTTR, A0A6J0AI81_VICPA, A0A1S3GPI0_DIPOR, Q568T8_DANRE. MEGA software (version 10.1.8) was used to do multiple sequence alignments and visualization.

## Statistical Analysis

All statistical analyses were conducted using GraphPad Prism 9.4 (GraphPad Software). All quantitative data were presented as the mean ± SD or SEM. Statistical analyses were performed with two-tailed Student’s *t*-test (when there were two groups), or one-way analyses of variance (ANOVA, when there were more than two groups)) or two-way ANOVA (when there were two or more groups with multiple subcategories within each group). *P* < 0.05 was considered statistically significant.

## Supplementary Materials for

**Extended Data Fig. 1.**
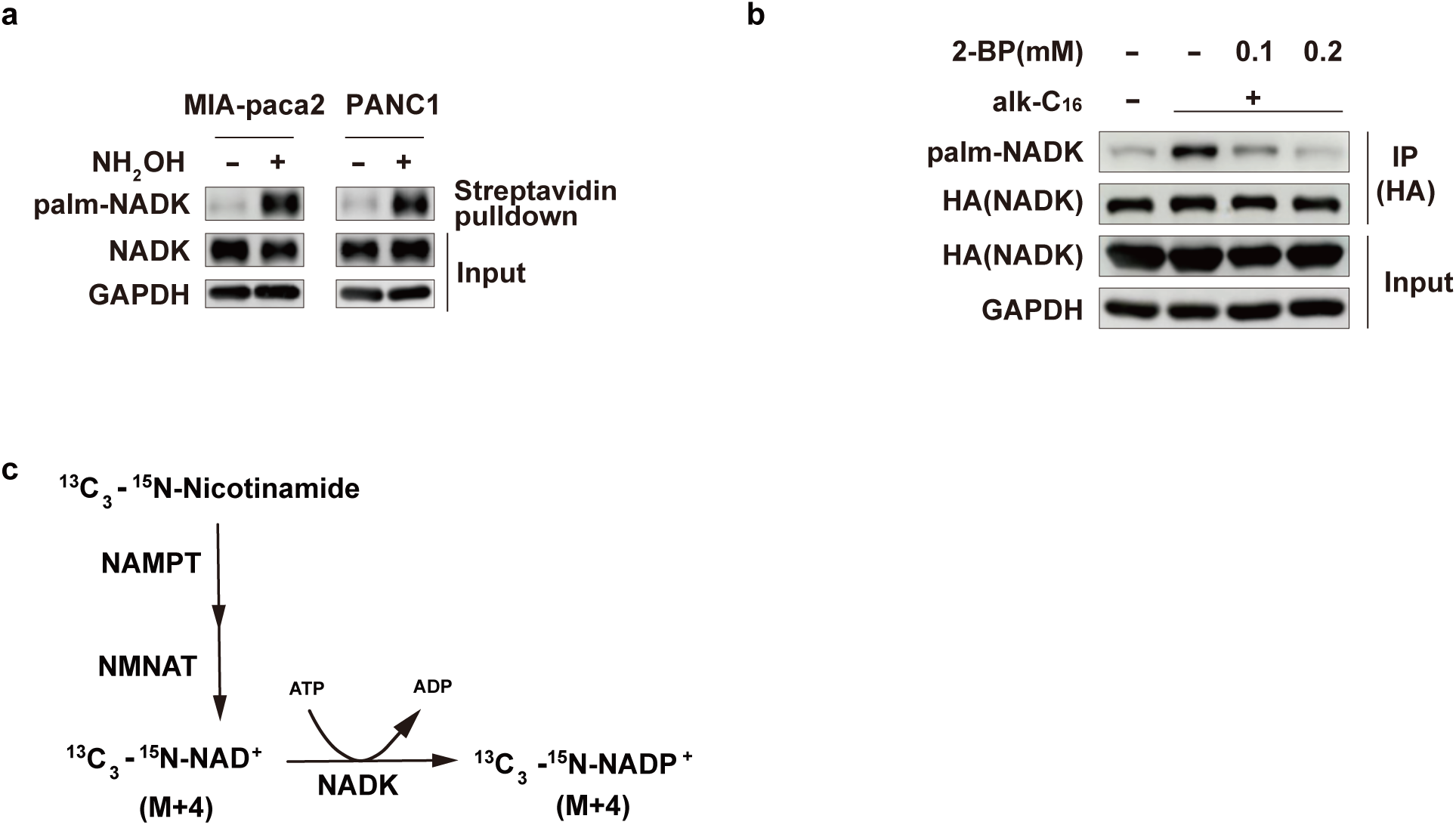
NADK is palmitoylated in cells. a. NADK palmitoylation was analyzed by ABE assay in MIA-paca2 and PANC-1 pancreatic cells. b. HEK293T cells transfected with HA-NADK were treated with 0 ∼ 0.2 mM 2-BP for 12 h, and metabolic labelled with 50 μM alk-C16 for 8 h, and NADK palmitoylation level was determined by CCR method. c. Schematic diagram of ^13^C3-^15^N-nicotinamide tracing for *in vivo* NADK function. NAMPT, nicotinamide phosphoribosyl transferase; NMNAT, nicotinamide mononucleotide adenylyl transferase.

**Extended Data Fig. 2.**
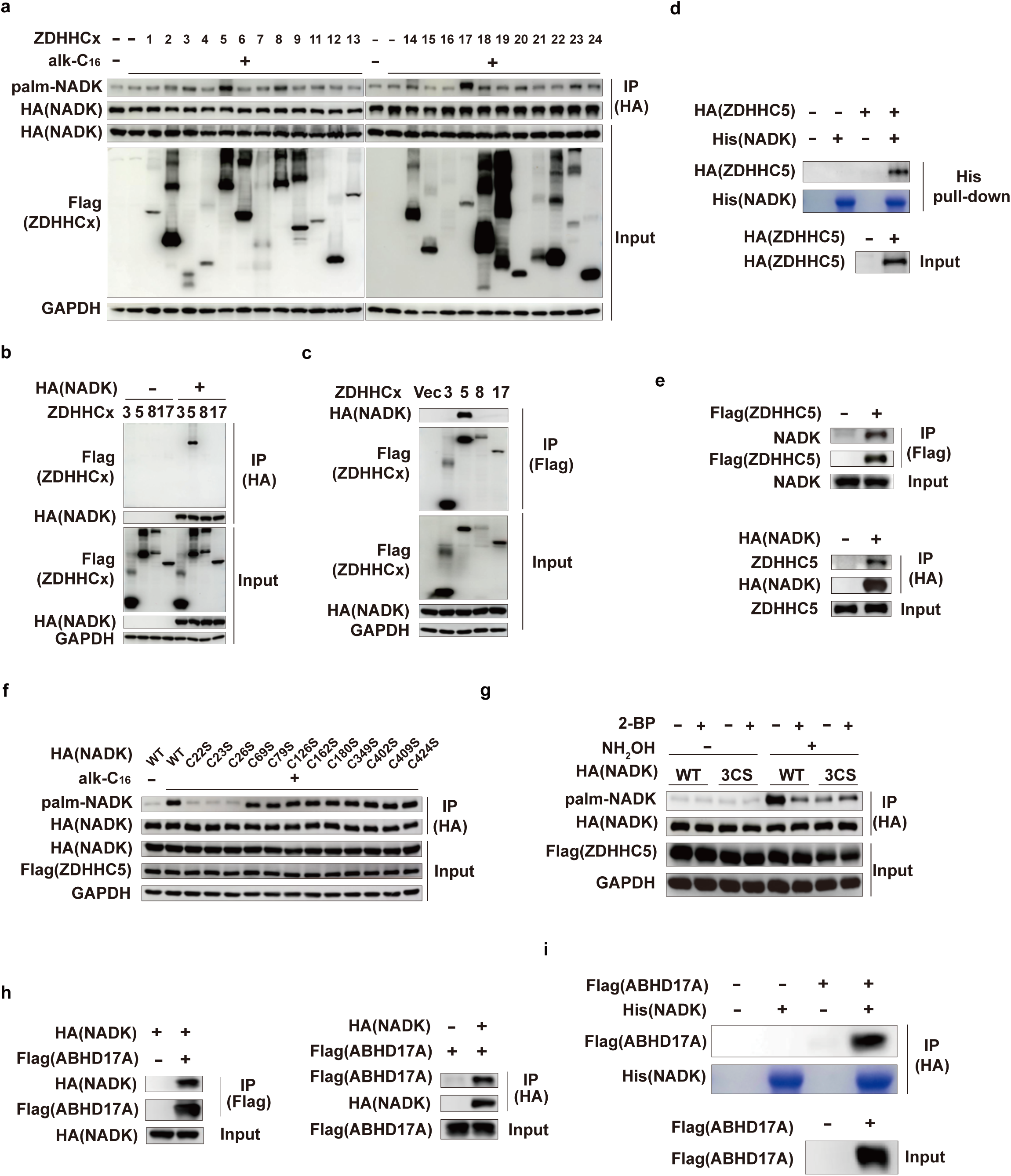
NADK is palmitoylated by ZDHHC5 at Cys22, 23 and 26. a. HEK293T cells were transfected with indicated plasmids and metabolic labelled with 50 μM alk-C16 for 8 h, and NADK palmitoylation level was determined by CCR method. b, c. Reciprocal co-IP assay for HA-NADK and selected ZDHHCx. d. His-Tag pull-down assays were performed to detect the direct interactions between NADK and ZDHHC5 *in vitro*. Purified His-NADK from *E. coli* bacteria pulled down the HA-ZDHHC5 expressed in HEK293T. e. HEK293T cells were transfected with the indicated plasmids, and the interaction between NADK and ZDHHC5 was verified by co-IP experiments. f. Endogenous NADK was depleted in HEK293T cells by sgRNA, followed by co-transfection with Flag-ZDHHC5 and HA-NADK WT or indicated cysteine mutants respectively. The cells were metabolic labelled with 50 μM alk-C16 for 8 h, and NADK palmitoylation level was determined by CCR method. g. Endogenous NADK was depleted in HEK293T cells by sgRNA, followed by stable reconstitution with cDNAs encoding HA-NADK WT or 3CS mutant. The cells were transfected with Flag-ZDHHC5, then treated with or without 2-BP (0.1 mM) for 12 h and NADK palmitoylation was analyzed by ABE assay. h. HEK293T cells were transfected with the indicated plasmids, and the interaction between NADK and ABHD17A was examined by reciprocal co-IP experiments. i. His-Tag pull-down assays were performed to detect the direct interaction between NADK and ABHD17A *in vitro*. His-NADK purified from *E. coli* bacteria pulled down the Flag-ABHD17A expressed in HEK293T.

**Extended Data Fig. 3.**
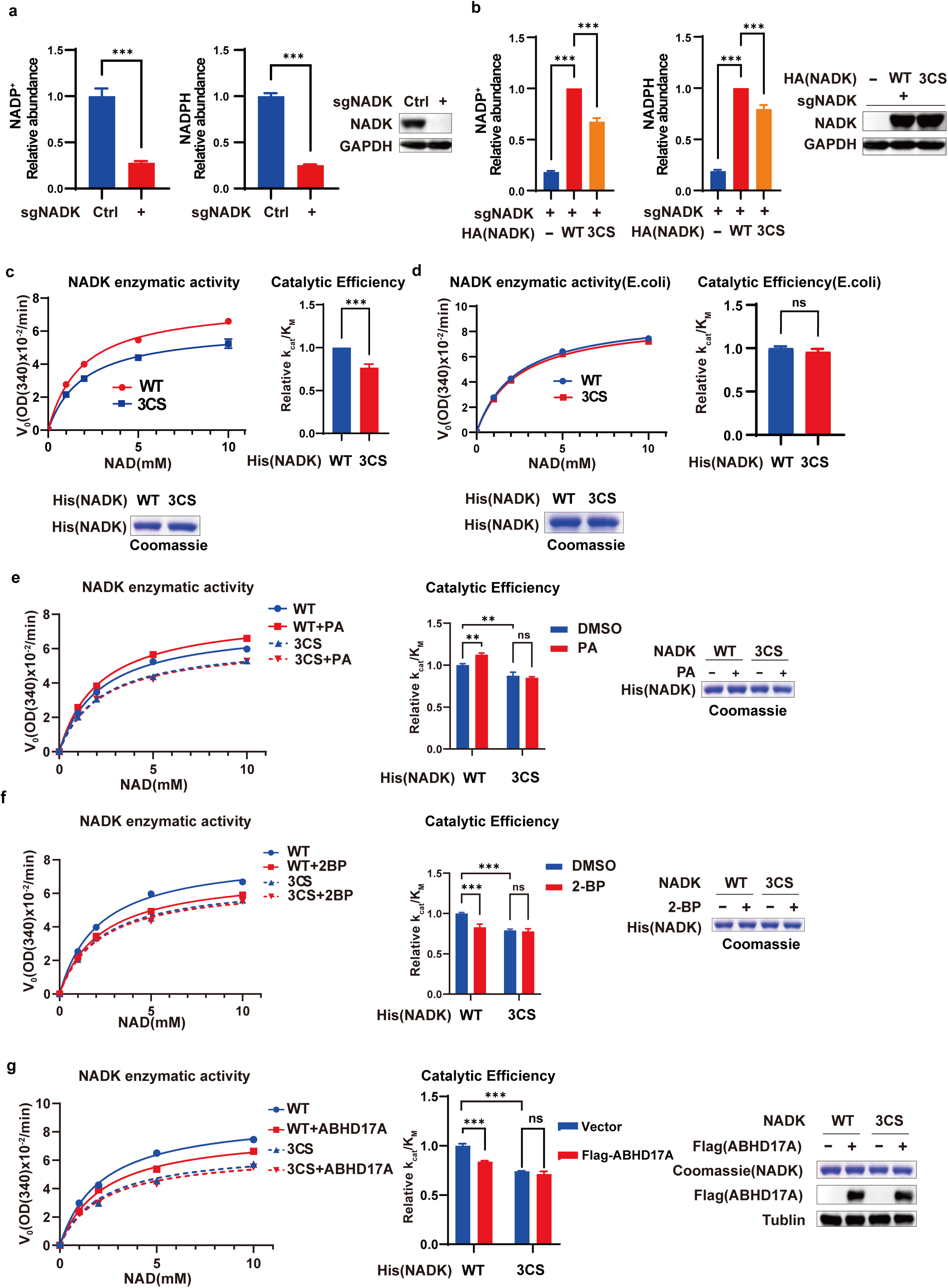
NADK palmitoylation boosted NADK activity. a. Endogenous NADK was depleted in HEK293T cells by sgRNA, relative abundance of NADP(H) were quantified by LC-MS. b. Endogenous NADK was depleted in HEK293T cells by sgRNA, followed by stable reconstitution with cDNAs encoding HA-NADK WT or 3CS mutant, relative abundance of NADP(H) were quantified by LC-MS. [a, b] Data are presented as the mean ± SD of biological duplicates and are representative of three independent experiments. [a] *P* values were determined by unpaired, two-tailed Student’s *t*-tests. [b] *P* values were determined by two-way ANOVA. \**P*<0.05, \*\**P*<0.01, \*\*\**P*<0.001. c. Endogenous NADK was depleted in HEK293T cells by sgRNA, followed by stable reconstitution with cDNAs encoding His-NADK-WT or 3CS mutant, and then His-NADK WT or 3CS mutant proteins were purified using Ni sepharose, and subjected to Coomassie staining and an NADK activity assay. d. Similar as in c., but the His-NADK WT or 3CS mutant proteins purified from *E. coli* were used instead. e, f. As in c, but the cells were treated with or without 0.1mM PA (e) or 2-BP (f) for 12h, and then his-NADK proteins were purified using Ni sepharose, and subjected to Coomassie staining quantification and an NADK enzymatic activity assay. g. As in c, but the cells were co-transfected with Flag-ABHD17A, and His-NADK proteins purification and NADK activity assay were performed and presented in the same way. [c to g] Data are presented as the mean ± SD of biological duplicates and are representative of three independent experiments. [c-d] *P* values were determined by unpaired, two-tailed Student’s *t*-tests. [e-g] *P* values were determined by two-way ANOVA. V0, reaction velocity; OD, optical density. \**P*<0.05, \*\**P*<0.01, \*\*\**P*<0.001.

**Extended Data Fig. 4.**
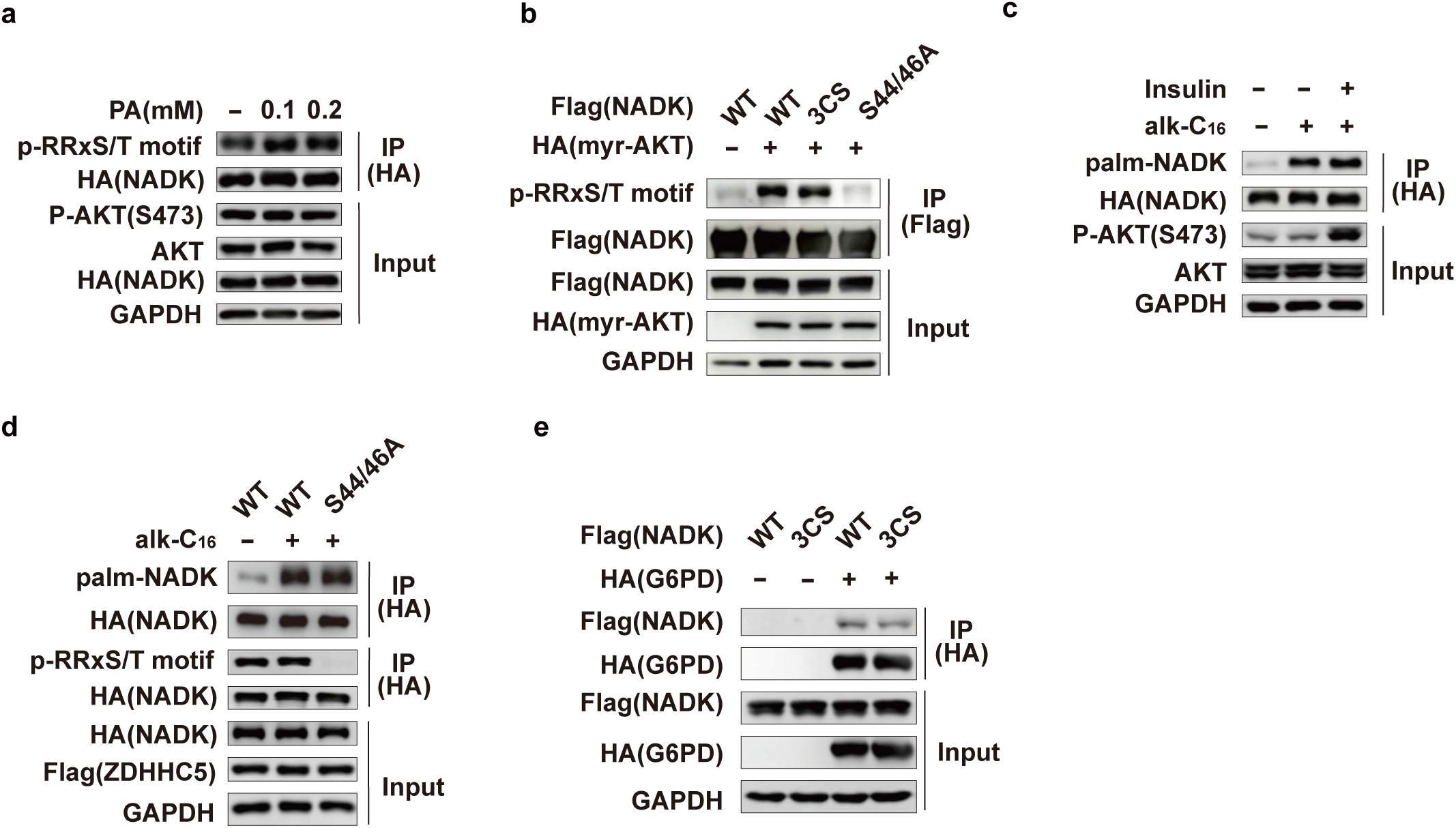
ZDHHC5-mediated NADK palmitoylation functions in parallel with Akt-mediated phosphorylation. a. HEK293T cells transfected with HA-NADK and AKT were treated with PA (0 ∼ 0.2 mM) for 12 h, then the immunoprecipitated NADK (with anti-HA affinity gel) were used to check phosphorylation. b. HEK293T cells transfected with the indicated plasmids, then the immunoprecipitated NADK (with anti-Flag affinity gel) were used to check phosphorylation. c. HEK293T cells transfected with the indicated plasmids were serum deprived for 12h, then stimulated with insulin (Ins) (0.5 mM, 2 h), and metabolic labelled with 50 μM alk-C16 for 8 h, and NADK palmitoylation level was determined by CCR method. d. HEK293T cells transfected with the indicated plasmids were metabolic labelled with 50 μM alk-C16 for 8 h, and NADK palmitoylation level was determined by CCR method. e. HEK293T cells were transfected with the indicated plasmids, and the interaction between NADK and G6PD was examined by co-IP experiments.

**Extended Data Fig. 5.**
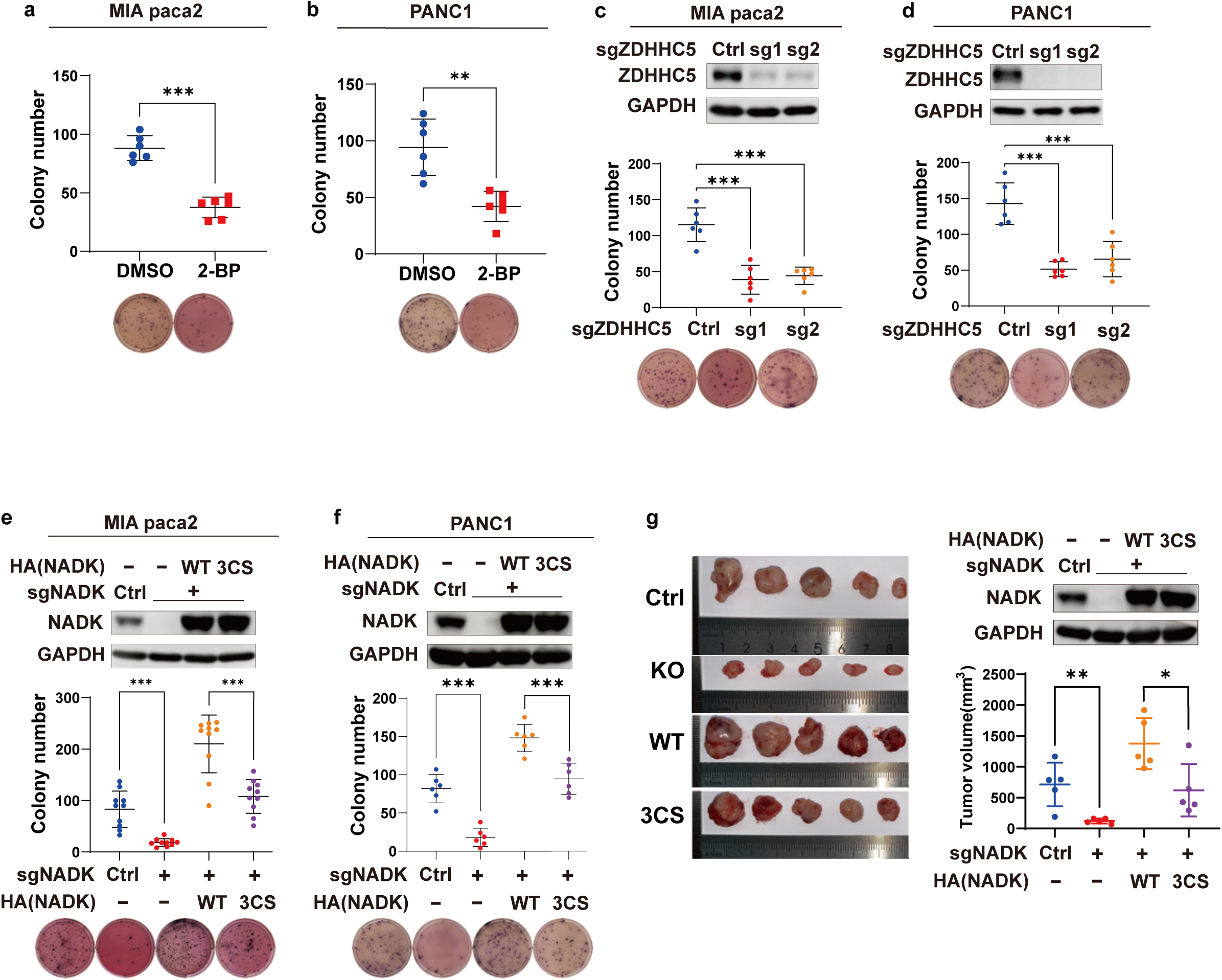
ZDHHC5 mediated NADK palmitoylation promotes tumor cell growth. a, b. Soft agar colony formation assay from MIA PaCa-2 (a) and PANC-1 (b) cells treated with 2-BP. n=6. Individual data and means ± SD are shown. *P* values were determined by unpaired, two-tailed *t*-test. c, d. Soft agar colony formation assay from MIA PaCa-2 (c) and PANC-1 (d) cells with stable sgRNA-mediated knockout of ZDHHC5. n=6. Individual data and means ± SD are shown. *P* values were determined by one-way ANOVA. e, f. Soft agar colony formation assay from MIA PaCa-2 (e) and PANC-1 (f) cells with stable sgRNA-mediated knockout of NADK and stably reconstituted with HA-NADK WT or 3CS mutant. MIA PaCa-2: n=10, PANC-1: n=6. Individual data and means ± SD are shown. *P* values were determined by unpaired, two-tailed *t*-test were used. g. NADK knockout PANC-1 cells stably reconstituted with HA-NADK WT or 3CS mutant or control vector were subcutaneously injected into athymic nude mice. Images of tumors at the end point of experiment, statistics on tumor volume, and NADK protein level are presented. *P* values were determined by unpaired, two-tailed Student’s *t*-tests. [a-g] \**P*<0.05, \*\**P*<0.01, \*\*\**P*<0.001.

**Extended Data Fig. 6.**
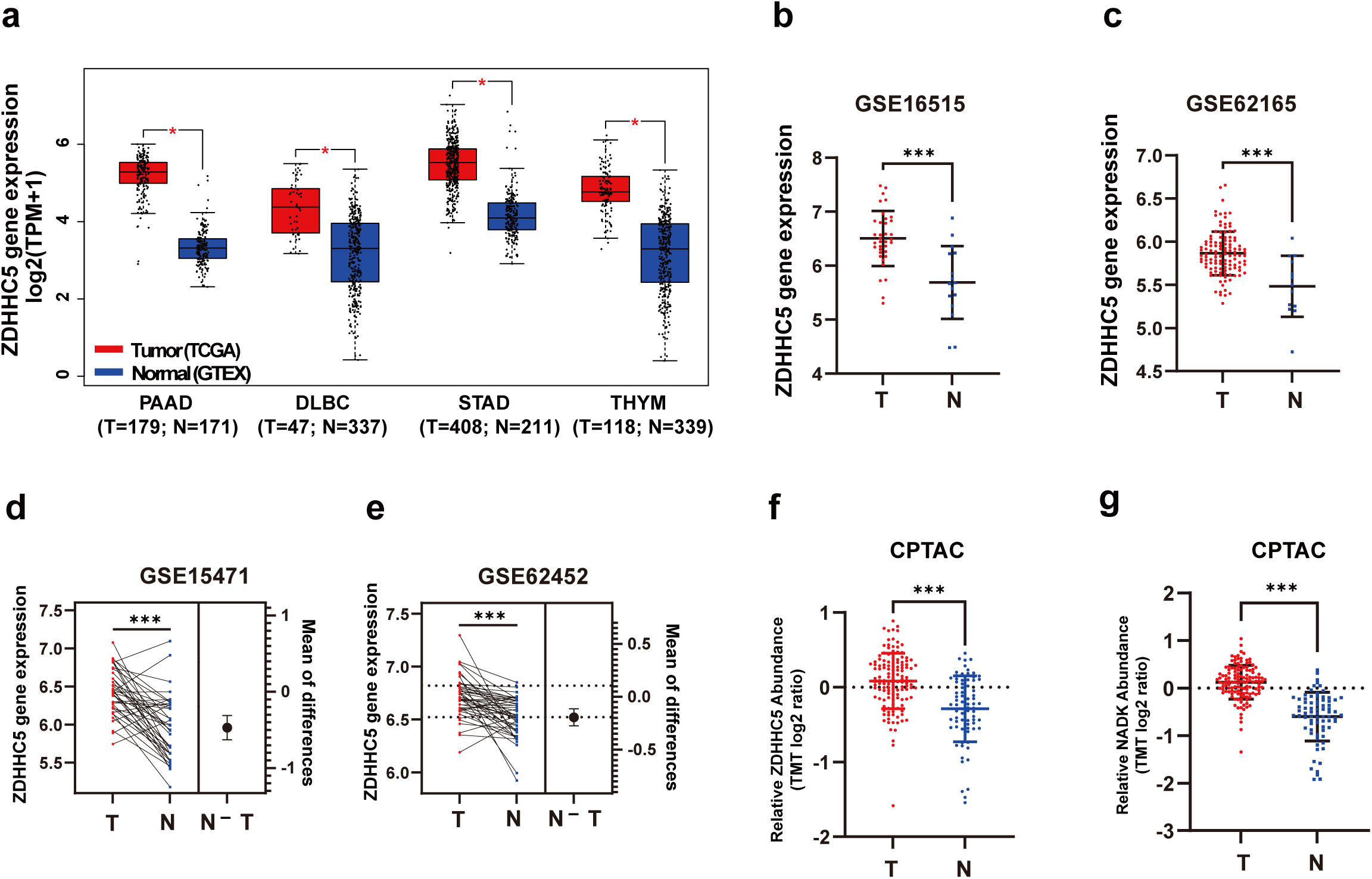
ZDHHC5 is elevated in human PDAC from the public database. a. Comparison of ZDHHC5 expression at mRNA levels in several tumor samples from the Cancer Genome Atlas (TCGA) database and normal controls from the Genotype-Tissue Expression (GTEx) database using GEPIA (http://gepia.cancer-pku.cn). Unpaired two-samples *t*-test was used, \**P*<0.01. b, c. ZDHHC5 expression at mRNA levels in PDAC tumor samples (T) and pancreatic normal controls (N) from the Gene Expression Omnibus (GEO) database (GSE16515 (b) and GSE62165 (c)). Data were obtained from GEO2R and plotted using GraphPad Prism 9.4. Unpaired, two-tailed *t-*test were used. \*\*\**P*<0.001. d,e. ZDHHC5 expression at mRNA levels in PDAC tumor (T) and adjacent pancreatic normal samples (N) from the GEO database (GSE15471 (d) and GSE62452 (e)). Data were obtained from GEO2R and plotted using GraphPad Prism 9.4. Paired two-tailed *t*-test were used. The right half of the graph shows the mean of difference between the two groups and the 95% confidence interval for the mean of difference. \*\*\**P*<0.001. f, g. ZDHHC5 (f) and NADK (g) expression at protein levels in PDAC tumor (T) and adjacent pancreatic normal samples (N) from the Clinical Proteomic Tumor Analysis Consortium (CPTAC) database (https://pdc.cancer.gov/pdc/). Data were obtained using cProSite (https://cprosite.ccr.cancer.gov/) and plotted using GraphPad Prism 9.4. Unpaired two-tailed *t*-test were used. \*\*\**P*<0.001.

**Extended Data Fig. 7.**
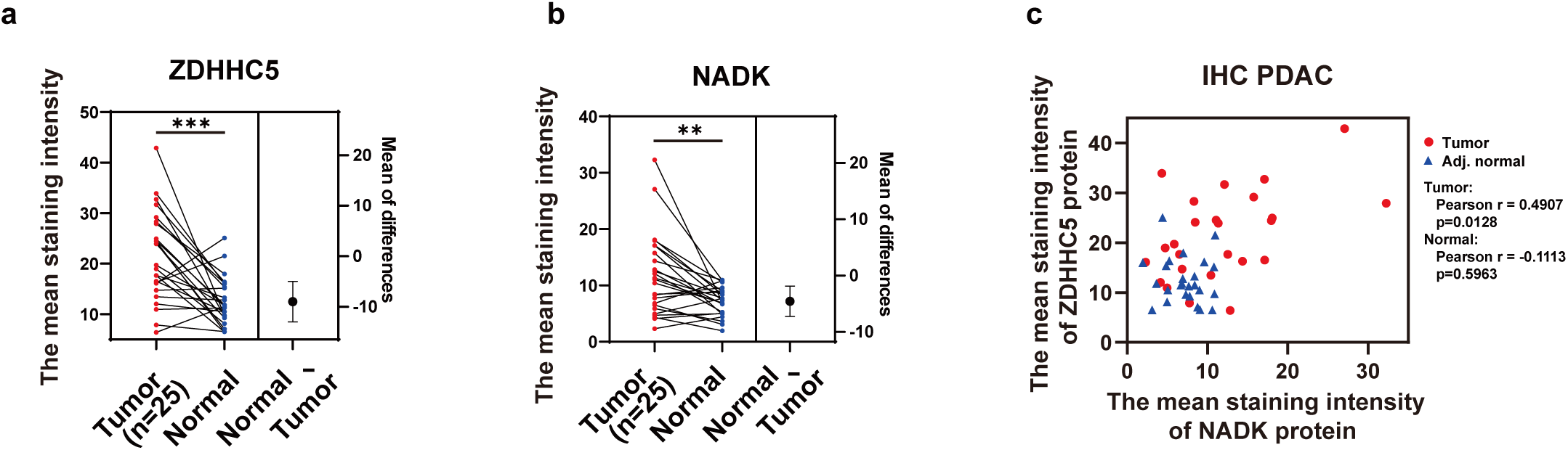
ZDHHC5 is upregulated in clinical PDAC samples. a. b. The mean staining intensity of ZDHHC5 (a) and NADK (b) in PDAC tumor and adjacent normal samples determined by IHC method and calculated using Image-Pro Plus software. Paired two-samples *t*-test were used. \*\*\**P*<0.001. c. Correlation between the mean staining intensity of ZDHHC5 and NADK in PDAC or pancreatic normal tissues. *P* values were calculated by two-tailed Pearson correlation. \*\**P*<0.01.

## References

1 Yap, L. P., Garcia, J. V., Han, D. & Cadenas, E. The energy-redox axis in aging and age-related neurodegeneration. Adv Drug Deliv Rev 61, 1283–1298, doi:10.1016/j.addr.2009.07.015 (2009).

2 Ju, H. Q., Lin, J. F., Tian, T., Xie, D. & Xu, R. H. NADPH homeostasis in cancer: functions, mechanisms and therapeutic implications. Signal Transduct Target Ther 5, 231, doi:10.1038/s41392-020-00326-0 (2020).

3 Jeon, S. M., Chandel, N. S. & Hay, N. AMPK regulates NADPH homeostasis to promote tumour cell survival during energy stress. Nature 485, 661–665, doi:10.1038/nature11066 (2012).

4 Lerner, F., Niere, M., Ludwig, A. & Ziegler, M. Structural and functional characterization of human NAD kinase. Biochemical and biophysical research communications 288, 69–74, doi:10.1006/bbrc.2001.5735 (2001).

5 Pollak, N., Niere, M. & Ziegler, M. NAD kinase levels control the NADPH concentration in human cells. J Biol Chem 282, 33562–33571, doi:10.1074/jbc.M704442200 (2007).

6 Shi, F., Li, Y., Li, Y. & Wang, X. Molecular properties, functions, and potential applications of NAD kinases. Acta Biochim Biophys Sin (Shanghai) 41, 352–361, doi:10.1093/abbs/gmp029 (2009).

7 Ishikawa, Y. & Kawai-Yamada, M. Physiological Significance of NAD Kinases in Cyanobacteria. Front Plant Sci 10, 847, doi:10.3389/fpls.2019.00847 (2019).

8 Love, N. R. et al. NAD kinase controls animal NADP biosynthesis and is modulated via evolutionarily divergent calmodulin-dependent mechanisms. Proceedings of the National Academy of Sciences of the United States of America 112, 1386–1391, doi:10.1073/pnas.1417290112 (2015).

9 Hoxhaj, G. et al. Direct stimulation of NADP(+) synthesis through Akt-mediated phosphorylation of NAD kinase. Science (New York, N.Y.) 363, 1088–1092, doi:10.1126/science.aau3903 (2019).

10 Hoxhaj, G. & Manning, B. D. The PI3K–AKT network at the interface of oncogenic signalling and cancer metabolism. Nature Reviews Cancer 20, 74–88, doi:10.1038/s41568-019-0216-7 (2020).

11 Rabani, R., Cossette, C., Graham, F. & Powell, W. S. Protein kinase C activates NAD kinase in human neutrophils. Free radical biology & medicine 161, 50–59, doi:10.1016/j.freeradbiomed.2020.09.022 (2020).

12 Schild, T. et al. NADK is activated by oncogenic signaling to sustain pancreatic ductal adenocarcinoma. Cell reports 35, 109238, doi:10.1016/j.celrep.2021.109238 (2021).

13 Martínez-Reyes, I. & Chandel, N. S. Cancer metabolism: looking forward. Nature reviews. Cancer 21, 669–680, doi:10.1038/s41568-021-00378-6 (2021).

14 Faubert, B., Solmonson, A. & DeBerardinis, R. J. Metabolic reprogramming and cancer progression. Science (New York, N.Y.) 368, doi:10.1126/science.aaw5473 (2020).

15 Munir, R., Lisec, J., Swinnen, J. V. & Zaidi, N. Lipid metabolism in cancer cells under metabolic stress. Br J Cancer 120, 1090–1098, doi:10.1038/s41416-019-0451-4 (2019).

16 Koundouros, N. & Poulogiannis, G. Reprogramming of fatty acid metabolism in cancer. Br J Cancer 122, 4–22, doi:10.1038/s41416-019-0650-z (2020).

17 Chamberlain, L. H. & Shipston, M. J. The physiology of protein S-acylation. Physiological reviews 95, 341–376, doi:10.1152/physrev.00032.2014 (2015).

18 Linder, M. E. & Deschenes, R. J. Palmitoylation: policing protein stability and traffic. Nature reviews. Molecular cell biology 8, 74–84, doi:10.1038/nrm2084 (2007).

19 Amendola, C. R. et al. KRAS4A directly regulates hexokinase 1. Nature 576, 482–486, doi:10.1038/s41586-019-1832-9 (2019).

20 Brownlee, C. & Heald, R. Importin α Partitioning to the Plasma Membrane Regulates Intracellular Scaling. Cell 176, 805–815.e808, doi:10.1016/j.cell.2018.12.001 (2019).

21 Lu, Y. et al. Palmitoylation of NOD1 and NOD2 is required for bacterial sensing. *Science (New York*, N.Y.) 366, 460–467, doi:10.1126/science.aau6391 (2019).

22 McCarthy, A. E., Yoshioka, C. & Mansoor, S. E. Full-Length P2X(7) Structures Reveal How Palmitoylation Prevents Channel Desensitization. Cell 179, 659–670.e613, doi:10.1016/j.cell.2019.09.017 (2019).

23 Ko, P. J. & Dixon, S. J. Protein palmitoylation and cancer. EMBO reports 19, doi:10.15252/embr.201846666 (2018).

24 Lin, H. Protein cysteine palmitoylation in immunity and inflammation. FEBS J 288, 7043–7059, doi:10.1111/febs.15728 (2021).

25 Lu, W., Wang, L., Chen, L., Hui, S. & Rabinowitz, J. D. Extraction and Quantitation of Nicotinamide Adenine Dinucleotide Redox Cofactors. Antioxid Redox Signal 28, 167–179, doi:10.1089/ars.2017.7014 (2018).

26 Qanbar, R. & Bouvier, M. Role of palmitoylation/depalmitoylation reactions in G-protein-coupled receptor function. Pharmacol Ther 97, 1–33, doi:10.1016/s0163-7258(02)00300-5 (2003).

27 Zhang, Y. et al. Upregulation of Antioxidant Capacity and Nucleotide Precursor Availability Suffices for Oncogenic Transformation. Cell metabolism 33, 94–109.e108, doi:10.1016/j.cmet.2020.10.002 (2021).

28 Zhang, Y. et al. G6PD-mediated increase in de novo NADP(+) biosynthesis promotes antioxidant defense and tumor metastasis. Sci Adv 8, eabo0404, doi:10.1126/sciadv.abo0404 (2022).

29 Tedeschi, P. M. et al. NAD+ Kinase as a Therapeutic Target in Cancer. Clin Cancer Res 22, 5189–5195, doi:10.1158/1078-0432.CCR-16-1129 (2016).

30 Rather, G. M., Pramono, A. A., Szekely, Z., Bertino, J. R. & Tedeschi, P. M. In cancer, all roads lead to NADPH. Pharmacol Ther, 107864, doi:10.1016/j.pharmthera.2021.107864 (2021).

31 Ilter, D. et al. NADK upregulation is an essential metabolic adaptation that enables breast cancer metastatic colonization. *bioRxiv*, 2022.2003.2025.485887, doi:10.1101/2022.03.25.485887 (2022).

32 Tsang, Y. H. et al. Functional annotation of rare gene aberration drivers of pancreatic cancer. Nature communications 7, 10500, doi:10.1038/ncomms10500 (2016).

33 Tang, Z. et al. GEPIA: a web server for cancer and normal gene expression profiling an d interactive analyses. Nucleic acids research 45, W98–W102, doi:10.1093/nar/gkx247 (2017).

34 Wang, D. et al. Abstract 3912: cProSite: A web based interactive platform for on-line proteomics and phosphoproteomics data analysis. Cancer research 82, 3912–3912, doi:10.1158/1538-7445.am2022-3912 (2022).

